# Hepatitis B Virus genomes associate with cellular sites of DNA damage by inducing replication stress

**DOI:** 10.1101/2024.03.21.586072

**Authors:** Gavin J. Marcoe, Clairine I. S. Larsen, Kinjal Majumder

## Abstract

Hepatitis B Virus (HBV) is a leading cause of liver cancer, with almost 300 million infected individuals worldwide. Although HBV-infected patients benefit from drug regimens that help to control chronic infection, they are rarely clinically cured of HBV. The HBV genome persists in the nucleus of infected hepatocytes in the form of a covalently closed circular DNA (cccDNA) molecule, a reservoir of HBV DNA molecules that serve as the template for reactivation of long-term chronic HBV. However, despite playing a central role in the viral life cycle, little is understood about where cccDNA molecules localize, why they are so stable, and how they impact the host nuclear compartment. Perhaps because of this, there are few treatments that target cccDNA, which is critical for eradication of clinical HBV. Here, we show that HBV infection induces a cellular DNA Damage Response (DDR) that is comparable with cells undergoing replication stress, and this cellular replication stress is initiated *after* the formation of cellular cccDNA molecules. Using a novel high-throughput chromosome conformation capture technology that monitors the localization of HBV cccDNA molecules, we show that cccDNA molecules persist in the vicinity of many cellular fragile sites. Induction of cellular DNA damage leads to relocalization of the viral HBx oncoprotein to DDR sites in an ATM, ATR and DNA-PK dependent manner. Our findings contribute to the understanding of how HBV cccDNA navigates the host nuclear environment, identifying functional targets for development of therapies against HBV infection and resulting liver cancer.

**Importance:** Hepatitis B Virus (HBV) is the leading infectious cause of liver cancer globally. The virus persists in the nucleus long term by forming reservoirs in human liver cells. We have discovered that the HBV DNA localizes to sites on the host genome associated with DNA damage, and in doing so, HBV interferes with the host’s ability to efficiently amplify itself. This results in the induction of cellular DNA breaks, which we propose contributes to eventual cancer progression. Our findings provide new insights into how HBV infection may lead to liver cancer.

## Introduction

Nearly 300 million people across the globe suffer from chronic Hepatitis B virus (HBV) infection. Every year, over 800,000 deaths can be attributed to the effects of chronic infection, such as hepatocellular carcinoma (liver cancer) or liver cirrhosis. A significant number of new infections are caused by vertical transmission, from mother to child during pregnancy or upon breastfeeding (1). Once infected, Hepatitis B virions enter hepatocytes via receptor-mediated endocytosis using the sodium taurocholate co-transporting polypeptide (NTCP) entry receptor. Once inside the cytoplasm, the viral genome in the form of relaxed circular DNA (rcDNA), undergoes nuclear import to enter the host cell’s nucleus (2). In the nucleus, rcDNA is converted into the covalently closed circular DNA (cccDNA) with the aid of host DNA repair proteins (3). This form of the viral genome plays a central role in the life cycle of the virus where cccDNA acts as a transcription template for the production of several RNA forms including precoreRNA which is translated into the Hepatitis B E-antigen [HBeAg; (4, 5)]. Unfortunately, little is known about where the cccDNA localizes to and how it utilizes host factors. Understanding these critical aspects of HBV cccDNA’s navigation of the nuclear environment will help to identify functional targets to fully eliminate HBV infection from the liver. While there is an HBV vaccine, antiviral therapies are not curative due to persistence of cccDNA in hepatocytes and lack of widely available drugs that target cccDNA molecules (6–8). Therefore, it is critical to understand where cccDNA molecules localize in the nuclear milieu, how they persist long-term and how cccDNA persistence alters the host environment.

Viruses provoke a cellular DNA damage response (DDR) in the host nuclear compartment, generated by viral genomes, transcripts and proteins that are expressed during infection. This cellular DDR is amplified by the phosphatidylinositol 3-kinase-related kinases ATM, ATR and DNA-PK, which can have pro-viral or anti-viral effects (9). As a result, viruses have evolved distinct strategies to usurp or inactivate host DDR signals. Tumor viruses like Human Papillomavirus (HPV), Polyomaviruses and Epstein-Barr Virus (EBV) are regulated by ATM/ATR signaling (10–17), whereas other DNA viruses like Adenoviruses and Herpes Simplex Virus (HSV) have a more complicated relationship with the host DDR signaling kinases (18–21). The single-stranded DNA parvovirus Adeno-Associated Virus (AAV) induces a DNA-PK-dependent cellular DDR signaling cascade (22), whereas its relative Minute Virus of Mice (MVM) induces an ATM-dependent DDR (23). Interestingly, the life cycle of HBV in host hepatocytes is dependent on pan-nuclear ATM and ATR signaling (24). Additionally, the HBV core protein HBc is a substrate for ATM phosphorylation, implicating ATM-mediated signaling events in regulating the HBV life cycle (25). Since cellular ATM and ATR signaling activate chromatin modifiers, downstream kinases, host polymerases and DNA repair pathways in the host nuclear environment (9), it remains unknown which of these pathways regulate HBV life cycle in the nucleus.

HBV genomes enter the host nuclear environment in the form of partially single- and partially double-stranded DNA molecules called relaxed circular DNA (rcDNA). Conversion of rcDNA into cccDNA is carried out by host replication and repair proteins such as DNA Polymerase Kappa, Flap Endonuclease 1 (FEN1) and host DNA ligases (7, 26–28). Upon formation, cccDNA molecules act as transcription templates, generating pregenomic RNA (pgRNA), messenger RNAs (mRNAs), and precore RNA. The pgRNA undergoes reverse transcription within newly created nucleocapsids to form more rcDNA molecules that generate progeny infectious particles in the cytoplasm (5). Expression of HBV genes produce HBx, HBc, E antigen (HBeAg), polymerase and the small (S), medium (M) and large (L) surface proteins in infected cells (29). Out of these viral factors, only the viral polymerase is known to associate with the viral origins to facilitate replication. Additionally, the HBV oncoprotein HBx interacts with host Structural Maintenance of Chromosomes (SMC) proteins SMC5/6 that are required for efficient cellular DDR leading to homologous recombination [HR; (30)]. Since HR signaling in the cell is regulated by ATM and ATR kinases, these findings suggest that HBV utilizes these pathways through HBx. Consistently, ATR-mediated signals have also been implicated in facilitating cccDNA formation (31).

One of the consequences of virus-induced DDR is the regulation of cell cycle entry or induction of cell cycle arrest. DNA tumor viruses such as HPV (32), Kaposi’s Sarcoma-associated Herpesvirus [KSHV; (33, 34)], Epstein-Barr virus [EBV; (35)], and Human T-Lymphotropic Virus 1 [HTLV1; (36)] antagonize cell cycle checkpoints to cause neoplastic transformation in infected cells. Paradoxically, while infection of primary human hepatocytes with HBV leads to G2/M arrest (37), infection of transformed HepG2 cells with HBV causes G1/S arrest (38). A likely unifying mechanism for HBV-induced cell cycle dysregulation is through the HBx protein that dysregulates cell-cycle checkpoint controls (39).

The cellular genome contains regions that preferentially accrue DNA damage called fragile sites, generated by replication stress [termed early replicating fragile sites; (40, 41)] or formation of secondary structures in late-replicating DNA regions [termed common fragile sites; (42–44)]. Both types of cellular fragile genomic regions are caused by or lead to transcription-replication conflicts. Cellular fragile sites are sites of localization of diverse DNA viruses, including HPV (45, 46), MVM (47) and EBV (48). Interestingly, HPV genomes are tethered to genomic fragile sites using host chromatin factors such as Bromodomain-containing protein 4 (BRD4) whereas EBV and MVM induce DNA damage at fragile genomic regions through unknown mechanisms (45, 49, 50). Prior studies investigating the localization of HBV genomes to cellular sites have discovered that transcriptionally inactive forms of HBV localize to distinct sites on the human genome, likely due to distinct chromatin signatures (51). Independently, HBV localization sites on the human genomes are enriched in binding sites of the cellular transcription factor Yin-Yang 1 [YY1; (6, 52)]. However, it remains unclear what form of the viral genome associates with these cellular sites. Additionally, the cause-effect relationship between host chromatin modification, transcription, and HBV cccDNA localization remains unknown.

In this study, we investigated the mechanisms by which HBV impacts host genome stability and associates with cellular DDR pathways. We show that HBV-induced replication stress is initiated immediately upon infection and exacerbated over time. As infection progresses, this replication stress leads to the induction of cellular DDR signals that colocalize with viral genomes and viral proteins. HBV cccDNA molecules associate with host fragile genomic regions, many of which are enriched in a distinct subset of cellular transcription factors, such as the C/EBP Homologous Protein (CHOP), known to cause liver cancer. Strikingly, focused induction of cellular DNA damage by laser micro-irradiation leads to relocalization of HBx to DDR sites in a PI3-kinase-like-kinase dependent manner. Our findings indicate a complex interplay between host cell genomic organization, DDR machinery, viral proteins, and viral genomes, with HBV cccDNA being preferentially and stably fixed to areas of basal DNA damage.

## Results

### HBV cccDNA molecules induce DDR signals in host cells

To determine how HBV cccDNA molecules impact host genome stability, we performed immunofluorescence analysis in HepG2-NTCP cells infected with HBV at 20 genome equivalents (GEQ) for 3 and 5 days (schematized in Fig. 1A). It has previously been shown that HBV rcDNA is converted into cccDNA at 3 days post-infection [3 dpi; (53, 54)]. To investigate the impact of HBV infection on cellular γH2AX levels at single-cell resolution, we performed confocal imaging for γH2AX foci in HBV-infected HepG2-NTCP cells that were co-stained for HBV core protein to identify infected cells. HBV-infected cells at 3 dpi had a median number of 0.5 γH2AX foci, which was comparable to that of Mock-infected cells at 3 days (Fig. 1B-C). However, at 5 dpi, HBV-infected cells had a median number of 6.0 γH2AX foci. Interestingly, it has previously been shown that cccDNA production is initiated at 3 dpi and plateaus at 5 dpi (53, 54). We monitored the impact of cccDNA molecule persistence on the cellular DDR by measuring the levels of phosphorylated H2AX (γH2AX) in whole-cell lysates at 5 dpi. As shown in Fig. S1, HBV infection led to an increase in total nuclear γH2AX at 5 dpi, which was slightly lower than that of HepG2-NTCP cells that had been treated with the DNA damaging agent hydroxyurea [HU; (55)]. This suggested that conversion of rcDNA to cccDNA is associated with induction of cellular DDR.

**Figure 1:**
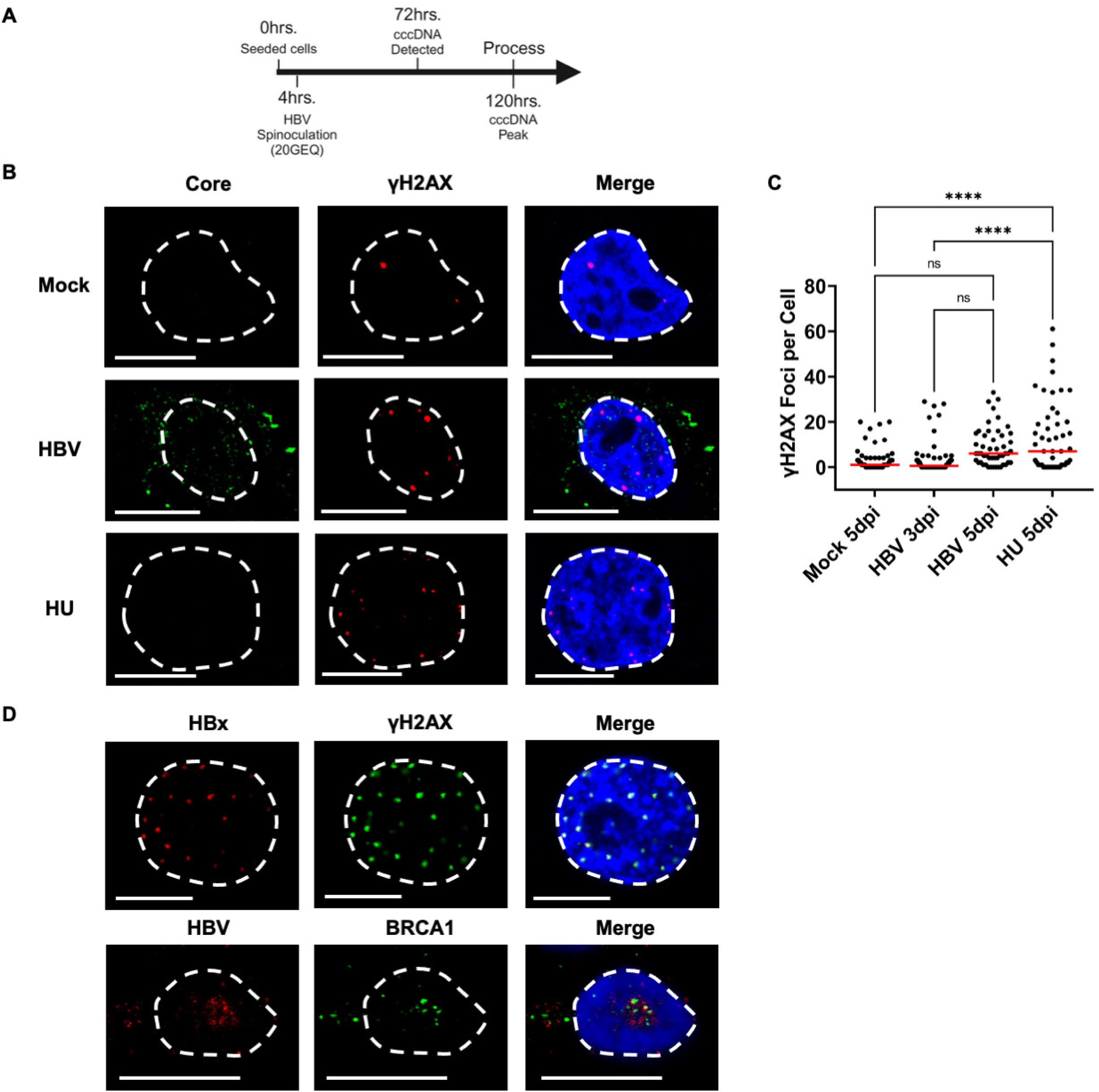
HBV cccDNA molecules induce DDR signals in host cells. (A) Schematic of plating of HepG2-NTCP cells, HBV infection by spinoculation and processing for DNA damage by immunofluorescence. (B) Representative images of DDR induction in HBV infected HepG2-NTCP cells monitored by staining for the assembled core protein (green) assessed by γH2AX staining (red). The nuclei are marked by DAPI staining (blue), nuclear borders demarcated by dashed white lines and scale bars in the representative images represent 10 micrometers. HepG2-NTCP cells pulsed with hydroxyurea (HU) for 12 hours prior to processing for immunofluorescence were used as positive control for γH2AX staining. Data is representative of three independent experiments of independent infections. (C) Nuclei in multiple HBV infected nuclei (presented in 1B) were measured during independent viral infections at 3 dpi and 5 dpi. Infected cells were identified by staining for HBV core protein, and the number of γH2AX foci were counted. The red bar represents median values with statistical analysis performed using one-way ANOVA, multiple comparisons test. Statistical significance is represented by *, p < 0.05 and ****, p < 0.00005. ns represents not significant difference in between the treatment conditions. (D) Representative immuno-FISH assays showing the location of the HBx and HBV genome (red) relative to that of cellular DDR markers like γH2AX (green, top panel) and BRCA1 (green, bottom panel) in HBV-infected HepG2-NTCP cells at 5 dpi. The nuclei are marked by DAPI staining (blue), nuclear borders are demarcated by dashed white lines and scale bars represent 10 micrometers.

To independently confirm whether HBV genomes and oncoproteins associate with cellular sites of DNA damage, we monitored the nuclear location of HBV factors relative to that of cellular DDR markers. As shown in Fig. 1D, HBx foci (top) colocalized with γH2AX and HBV genomes (bottom) colocalized with BRCA1 signals in the nucleus. Taken together, these results suggest that HBV genomes and proteins contribute to the induction of cellular DDR signals at late timepoints of infection, localizing to the same subnuclear compartments.

### HBV infection induces replication stress on the host genome

Since DNA viruses induce cellular replication stress which precedes DDR signals, we predicted that HBV-mediated host replication stress may contribute to genome instability. As cellular replication stress activates the ATR pathway (9), we visualized the location of HBV genomes relative to that of ATR and replication stress factors in the nucleus of HepG2-NTCP cells. At 5 dpi, HBV genomes associated with cellular RPA70 (Fig. 2A, top), ATRIP (Fig. 2A, middle) and PCNA (Fig. 2A, bottom). We therefore examined the impact of HBV infection on host replication forks using single-molecule DNA Fiber Assay [DFA, schematized in Fig. 2B; (56)]. Briefly, DFA utilizes sequential pulsing of BrdU analogs chlorodeoxyuridine (CldU) and iododeoxyuridine (IdU) to measure how cellular replication forks are impacted upon introduction of genotoxic stress. HBV infection at 5 dpi led to shortening of cellular replication forks measured by IdU (Fig. 2D) and CldU (Fig. 2E). The median length of IdU-labelled cellular tracks decreased from 3.42 μm in mock-infected cells to 2.32 μm in the presence of HBV, while the CldU-labelled tracks decreased from 5.40 μm in mock cells to 3.90 μm in HBV-infected cells. Categorization of the host DNA fibers revealed an increase of forks containing new origin firings and decrease in progressing replication forks in HBV-infected cells compared with uninfected cells (Fig. 2F). These findings suggested that HBV infection may induce aberrant firing of new origins in the host cell.

**Figure 2:**
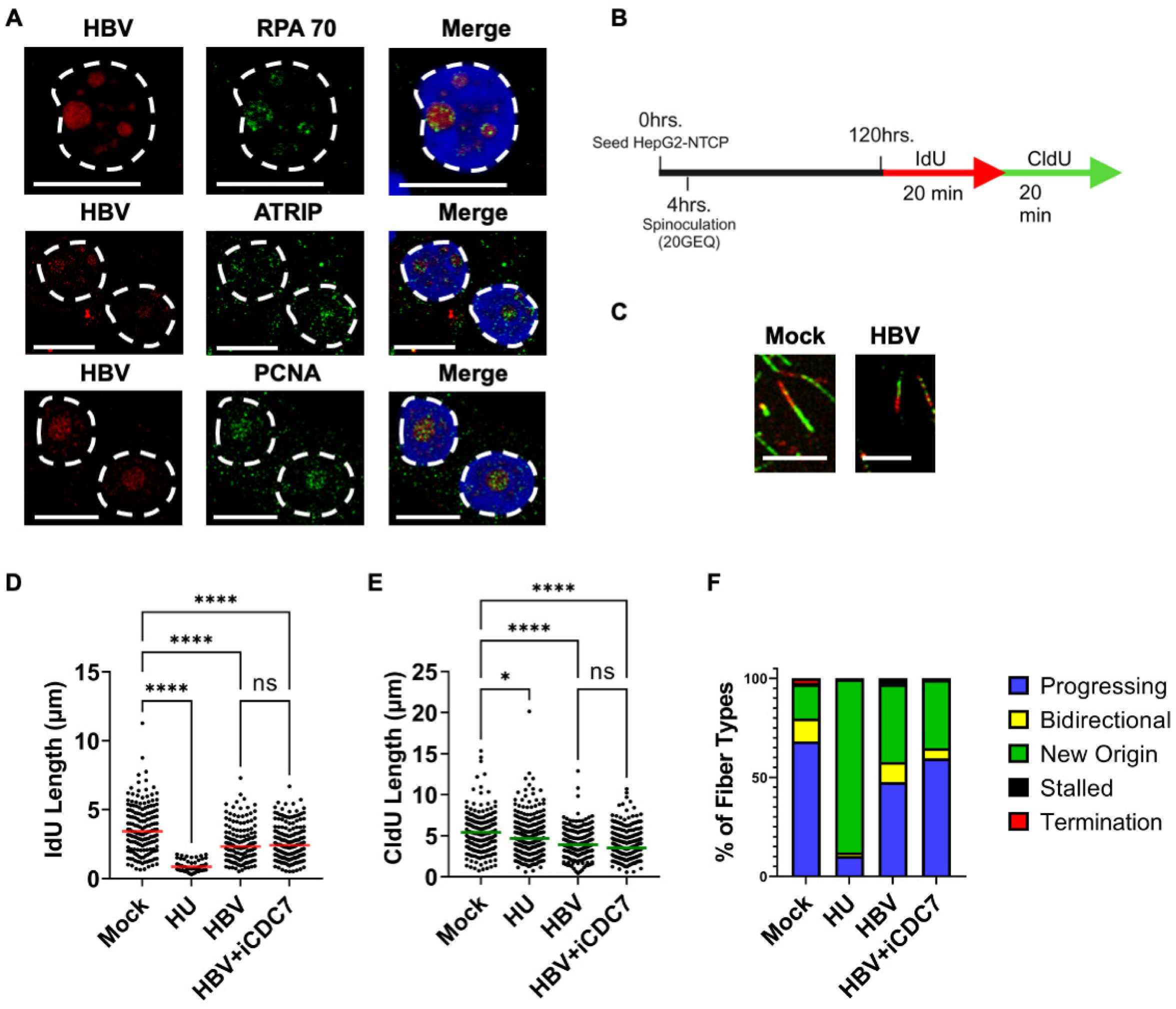
HBV infection induces replication stress on the host genome. (A) Representative immuno-FISH assays showing the location of the HBV genome (red) relative to that of cellular replication and replication stress markers like RPA70 (green, top panel), ATRIP (green, middle panel) and PCNA (green, bottom panel) in HBV-infected HepG2-NTCP cells at 5 dpi. The nuclei are marked by DAPI staining (blue), nuclear borders are demarcated by dashed white lines and scale represent 10 micrometers. (B) Schematic of HBV infection of HepG2-NTCP cells followed by sequential pulses of IdU and CldU prior to processing for DNA fiber analysis. (C) Representative fibers of HBV infected HepG2-NTCP cells at 5 dpi showing IdU (red) and CldU (green) incorporation. Scale bars represent 5 micrometers. (D,E) Individual fiber lengths were measured by quantifying the IdU and CldU lengths in HBV infected cells. Each datapoint represents the length of a single IdU and/or CldU labelled DNA fiber in HepG2-NTCP cells. The experiment was performed as described in the schematic described in 2B. At least 150 individual DNA fibers were measured for each condition across at least 2 independent infections. Similar results were obtained for at least two independent biological experiments of HBV infection. The horizontal lines (red for 2D and green for 2E) represent median values of all datapoints. Statistical analysis was performed using One Way ANOVA, multiple comparisons test, with **** representing p < 0.0005 and * representing p < 0.05. (F) Categorization of DNA fiber types as percentages of total of 100% as determined by presence of IdU or CldU that were divided into percentages that are progressing, bidirectional, stalled, terminated replication forks and new origin firings in HBV infected HepG2-NTCP cells.

Replication stress on the eukaryotic genome is regulated by phosphorylation of the Minichromosome Maintenance (MCM) helicase complex (57). Interestingly, inhibition of MCM helicase phosphorylation using the CDC7 inhibitor PHA767491 partially decreased new-origin firing (Fig. 2F) without impacting the median shortening of replication forks in HBV-infected cells (Fig. 2D, 2E), suggesting that HBV-induced replication stress is independent of MCM helicase activity. Taken together, these data show that HBV infection leads to induction of cellular replication stress, concurrent with the DDR induction that we observed using immunofluorescence analysis (Fig. 1B-C).

### HBV-induced replication stress increases as infection progresses

The correlation between HBV-induced cellular DDR and onset of cellular replication stress led us to ask whether replication stress is induced upon immediate entry of HBV viral particles into the nuclear compartment. To determine whether HBV infection leads to acute induction of cellular replication stress, we performed DFA at early timepoints post-infection (Fig. 3A-D). There was an immediate induction of replication stress at 1 hour post-infection (hpi), which persisted at the same level for four days (Fig. 3B,C). Characterization of replication events at these timepoints revealed an increase of new origin firings upon HBV infection, which increased further at 48 hpi, but disappeared by 96 hpi (Fig. 3D). Next, we asked whether the plasmid form of the viral genome was sufficient to induce replication stress. We transiently transfected HepG2 cells with the HBV-encoding plasmid TMA153 [hereafter referred to as pHBV; (58)] and performed DFA 24 hours post-transfection (Fig. 3E-H). When compared to cells transfected with an empty vector control, pHBV did not induce cellular replication stress (Fig. 3F-G) or alter the fractions of replication events (Fig. 3H), suggesting that the HBV genome itself is insufficient to induce cellular replication stress, instead requiring infection. Since HBV protein production is dependent on generation of transcribed viral RNA, these results further suggest that HBV pgRNA is insufficient to generate cellular replication stress. Interestingly however, transfection with pHBV lacking the ability to express HBeAg led to shortening of cellular replication forks, indicating (Fig. 3F-G) that HBeAg functions as a protective element that helps maintain the fidelity of host replication forks. Since HBeAg has previously been shown to decrease host p53 activity (59), we infer that intact p53 activity leads to increased replication fork processivity (60) in cells overexpressing HBV proteins, which manifests as shortened replication fibers. Together, these findings suggested that both infection and viral replication are required to induce and exacerbate cellular replication stress that leads to the induction of the downstream DDR.

**Figure 3:**
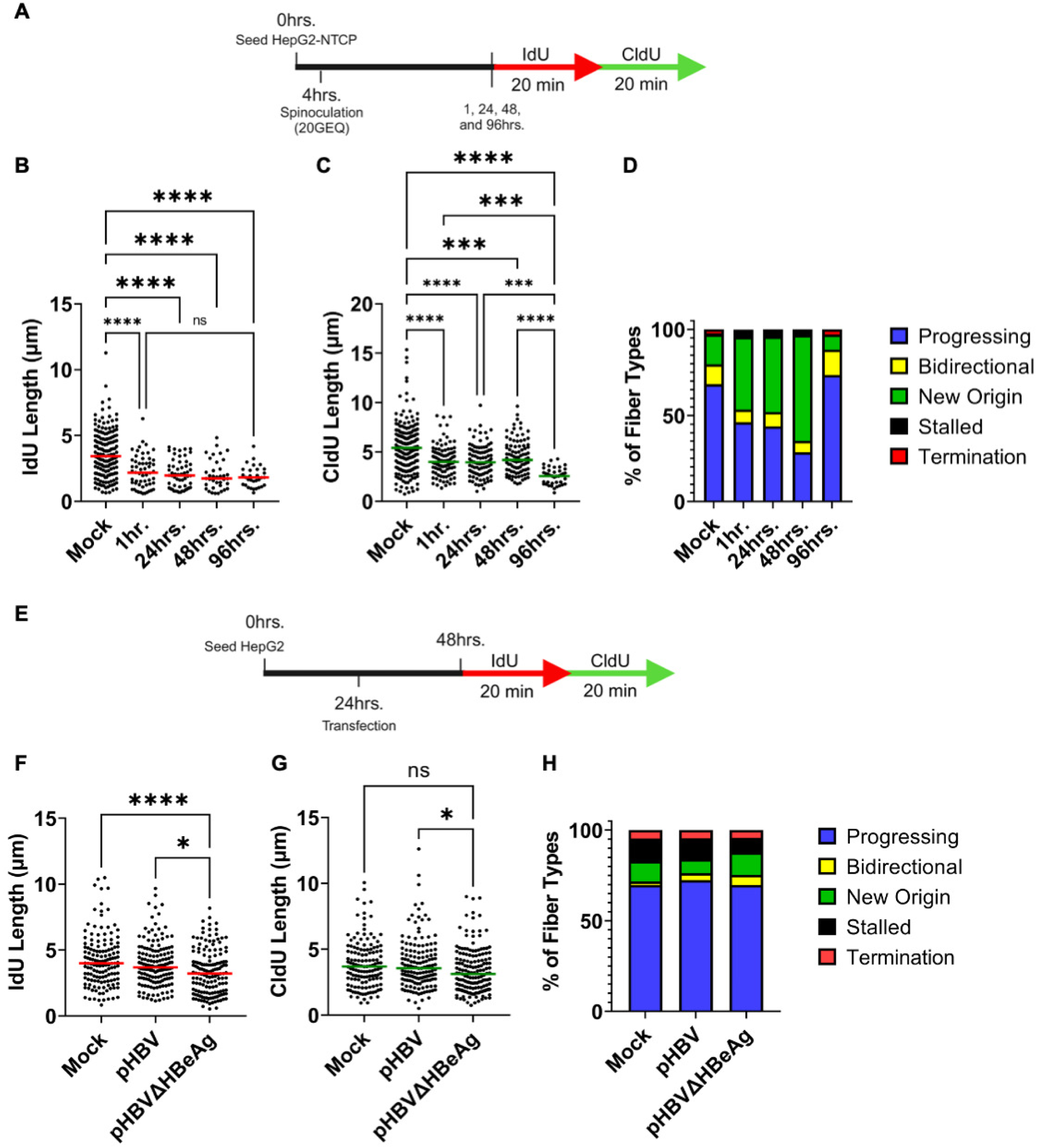
HBV-induced replication stress increases as infection progresses. (A) Schematic of HBV infection of HepG2-NTCP cells at different timepoints followed by IdU/CldU pulsing for 20 minutes each before processing for DNA fiber analysis. (B,C) Individual fiber lengths were measured by quantifying the IdU and CldU lengths in HBV infected cells at the indicated timepoints post-infection. Each datapoint represents the length of a single IdU and/or CldU labelled DNA fiber in HepG2-NTCP cells. The experiment was performed as described in the schematic described in Fig. 3A. At least 150 individual DNA fibers were measured for each condition across at least 2 independent infections. The horizontal lines (red for 3B and green for 3C) represent median values of all datapoints. Statistical analysis was performed using One Way ANOVA, multiple comparisons test, with **** representing p < 0.0005 and * representing p < 0.05. (D) Categorization of DNA fiber types as percentages of total of 100% as determined by presence of IdU or CldU that were divided into percentages that are progressing, bidirectional, stalled, terminated replication forks and new origin firings in HBV infected HepG2-NTCP cells at the indicated timepoints post-infection (56, 57). (E) Schematic of transfection of HBV infectious clone plasmids in HepG2 cells for 24 hours prior to IdU/CldU pulsing for DNA fiber analysis. (F,G) Individual fiber lengths were measured by quantifying the IdU and CldU lengths in HBV infected cells at 24 hours post-transfection. Each datapoint represents the length of a single IdU and/or CldU labelled DNA fiber in transfected HepG2 cells. The experiment was performed as described in the schematic described in 3E. At least 150 individual DNA fibers were measured for each condition across at least 2 independent infections. Similar results were obtained for at least two independent biological experiments of HBV infection. The horizontal lines (red for 3F and green for 3G) represent median values of all datapoints. Statistical analysis was performed using One Way ANOVA, multiple comparisons test, with **** representing p < 0.0005 and * representing p < 0.05. (H) Categorization of DNA fiber types as percentages of total of 100% as determined by presence of IdU or CldU that were divided into percentages that are progressing, bidirectional, stalled, terminated replication forks and new origin firings in HBV infectious clone transfected HepG2 cells at 24 hours post-infection.

### HBV cccDNA molecules associate with cellular fragile genomic regions

The development of genomics technologies, particularly high-throughput chromosome conformation capture assays, have provided essential tools that facilitated unbiased interrogation of viral genome association with cellular genomic sites (61, 62). These technologies have been previously deployed to track HBV genome localization, finding that HBV episomes associate with distinct genomic sites (51). However, the generation of multiple extrachromosomal forms of the viral genome during replication in the host, such as rcDNA, double-stranded linear DNA (dslDNA) and cccDNA (54), make it challenging to track the genomic forms associated with the host. To specifically identify sites of association of cccDNA molecules with the host genome, we have modified the chromosome conformation capture assay protocol, incorporating a T5 exonuclease treatment that degrades rcDNA and dslDNA (Fig. 4A). This technique, which we dub V3C-T5-seq, has enabled us to track the nuclear localization of cccDNA relative to the host genome. As shown in the representative genome browser track for chromosome 10 in Fig. 4B, the cccDNA molecules (bottom track) associate with similar genomic sites as all HBV genomic forms (top track). We validated that HBV genomes associate with the 10p12.3 site using fluorescence in situ hybridization on three-dimensionally preserved nuclei (3D-FISH, Fig. 4C). Importantly, we observed HBV genomes associated with all chromosomes except Chr 12, 13, 14, 18, 20, 21 and 22 (Fig. S2). Whole genome comparison of episomal HBV molecules with that of cccDNA-associated sites revealed that 44% of all HBV-associated genomic sites are made up of cccDNA molecules (Fig. 4D; statistical significance of the correlation is shown in Fig. 4E). These correlations are significant because the cccDNA molecules make up a small fraction of HBV episomes. Thus, the specificity and commonality of these cccDNA associations might indicate that these molecules are tethered to the host genome.

**Figure 4:**
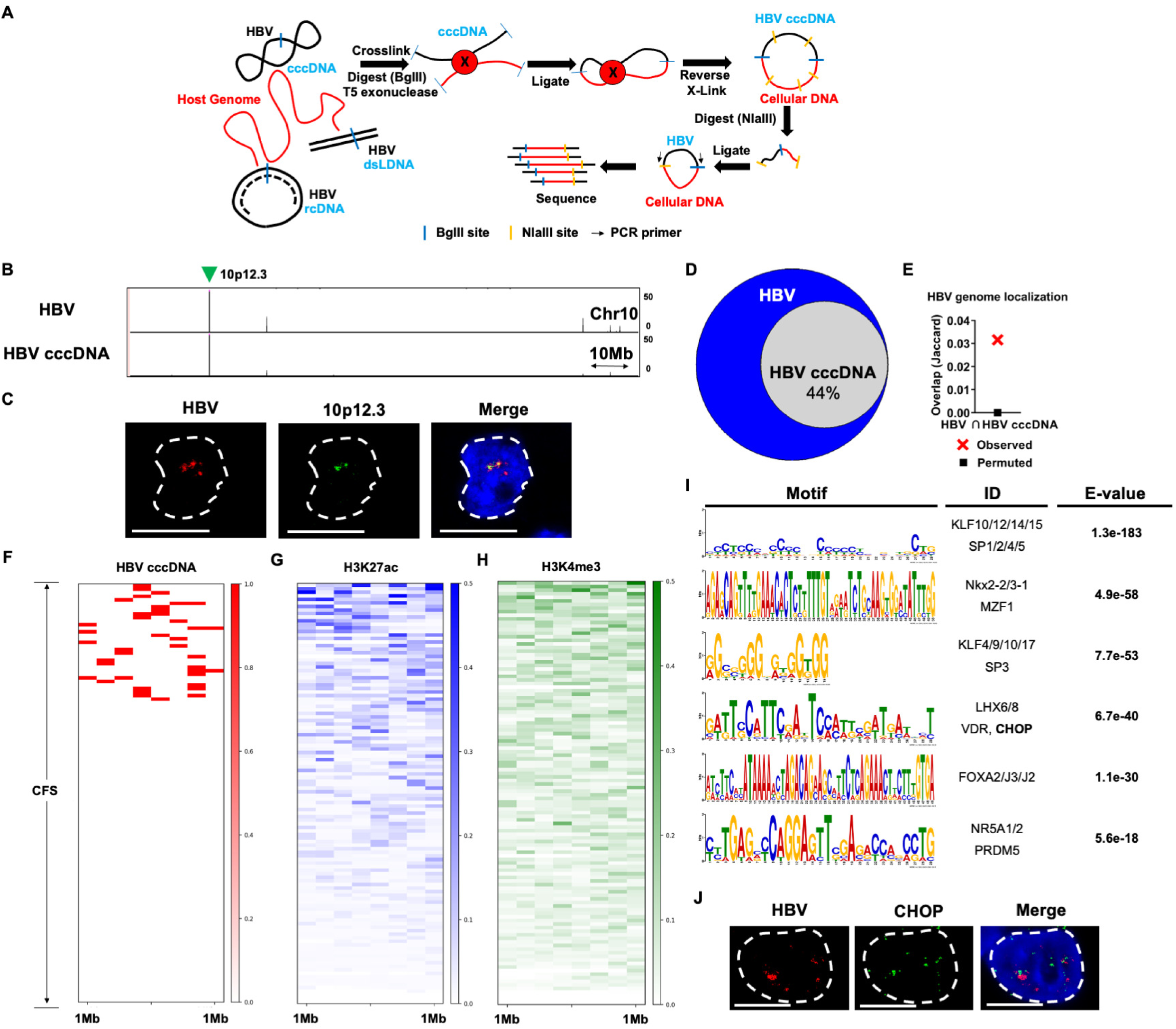
HBV cccDNA genomes associate with cellular fragile genomic regions. (A) Schematic of viral chromosome conformation capture assay that incorporates T5 Exonuclease treatment to capture the localization sites of HBV cccDNA molecules on the human genome in HepG2-NTCP cells. (B) Representative UCSC genome browser plots on human chromosome 10 comparing where all HBV genome forms localize (top) relative to that of HBV cccDNA molecules (bottom). The x-axis represents distance along the indicated human chromosome and y axis represents sequencing reads. The location of the 10p12.3 peak used for 3D-FISH validation is indicated by a green arrow over the track. (C) Validation of HBV genome localization to the indicated 10p12.3 peak on the human genome using representative FISH assay. The HBV genome is labelled in red, 10p12.3 region in green and the nucleus by DAPI staining. Nuclear borders are demarcated by white dashes, and the white line horizontal line represents a scalebar of 10 micrometers. (D) Venn diagrams comparing the total genomic regions associated with all HBV genomic forms (blue circle) compared with those sites that are associated with HBV cccDNA molecules (overlap indicated by gray circle). The cccDNA-associated sites make up 44% of all HBV localization sites. (E) Statistical analysis of total HBV localization relative to that of cccDNA molecules using Jaccard analysis where the intersection is represented by red crosses (Observed). The control intersection was calculated by Jaccard analysis where cccDNA-associated sites were intersected with a randomly generated library of genomic sites of the same size and number as total HBV localization, which is indicated by a black square (Permuted). Jaccard analysis values range from 0 to 1, with 0 indicating no intersection and 1 indicating complete intersection. (F-H) The locations of cellular fragile sites were compared relative to that of (F) HBV cccDNA localization, (G) transcriptionally active H3K27ac chromatin and (H) transcriptionally active H3K4me3 chromatin in HepG2 cells using Deeptools program on the Galaxy project analysis platform. (I) MEME and TOMTOM analysis platforms were used to compute the over-represented motifs and their corresponding transcription factor binding sites on the human genome that are associated with cccDNA molecules. The statistical significance of the identified transcription factors is represented by the respective E values. (J) Immuno-FISH validation of HBV genome (red) relative to that of the cellular transcription factor CHOP (green) was visualized in HBV infected HepG2-NTCP cells at 5 dpi. The nuclei were visualized by DAPI staining, nuclear borders are demarcated by white dashed line and the horizontal scale bar represents 5 micrometers.

Since small DNA viruses have been found associated with cellular sites of DNA damage (45, 47), many of which are fragile genomic regions (40–42, 45, 46), we compared the cccDNA-associated sites genome wide with that of previously identified cellular common fragile sites [CFS;(63)]. Cellular DNA damage forms multi-megabase (Mb) platforms of γH2AX on either side of DNA break sites (64). Therefore, we selected 1 Mb windows on either side of CFSs for comparative analysis and found that approximately one-third of CFSs were associated with cccDNA molecules within their 1 Mb vicinity (Fig. 4F). Furthermore, we observed that most fragile sites are associated with active cellular chromatin in HepG2 cells, monitored by H3K27 acetylation [Fig. 4G; (65)] and H3K4 tri-methylation [Fig. 4H; (65)], indicating that cccDNA molecules localize to a transcriptionally active subset of cellular fragile genomic sites. Since cellular fragile genomic regions are often caused by transcription-replication conflicts (66), we performed in-silico analysis using the MEME and TOMTOM bioinformatics suites to determine which host transcription factor consensus sequences might be upregulated in the cccDNA localization sites (67). These localization motifs are represented in Fig. 4I and the statistical significance of these findings is represented in the right column. These consensus sequences were associated with host-cell transcription factor binding sites, indicated in the middle column (Fig. 4I). Several of these host factors are associated with varying liver cancers and tumors, including Nkx2-2, MZF1, LHX6, LHX8 and VDR. Among these factors, CHOP has previously been shown to be associated with HBV-induced liver cancers (68). Excitingly, CHOP colocalized with the HBV genome in HBV-infected HepG2-NTCP cells at 5 dpi (Fig. 4J) establishing proof-of-concept for our analytical pipeline. Taken together, these findings validate the localization of HBV genomes to cellular sites enriched in CHOP binding.

### HBx proteins localize to cellular DNA breaks using host DDR kinase signaling

Because we observed induction of cellular replication stress and DDR response upon HBV infection as well as localization of HBV episomes to sites of DNA damage, we next wished to determine the response of HBV proteins and genomes to artificially induced DNA damage. We have previously used laser micro-irradiation to monitor how parvovirus genomes and non-structural proteins localize to cellular sites of DNA damage (69), so we utilized this approach coupled with immunofluorescence analysis in HepG2-NTCP cells infected with HBV for 5 days. As shown in Fig. 1D, HBx colocalized with cellular DDR sites marked with γH2AX. Upon induction of cellular DDR using laser micro-irradiation at distinct nuclear fields of view (monitored by γH2AX staining), HBx relocalized to the newly formed γH2AX sites (Fig. 5A, Mock panel; Fig. 5B, green line). To determine which host signals drive relocalization, we treated the infected HepG2-NTCP cells with inhibitors to key PI3-kinase-like kinases (Fig. 5A-B). Treatment of cells with ATM, ATR and DNA-PK inhibitors individually and in combination led to attenuated HBx relocalization to γH2AX sites. These observations indicate that HBx interaction with cellular DDR sites is dependent on signaling by PI3-kinase-like kinases.

**Figure 5:**
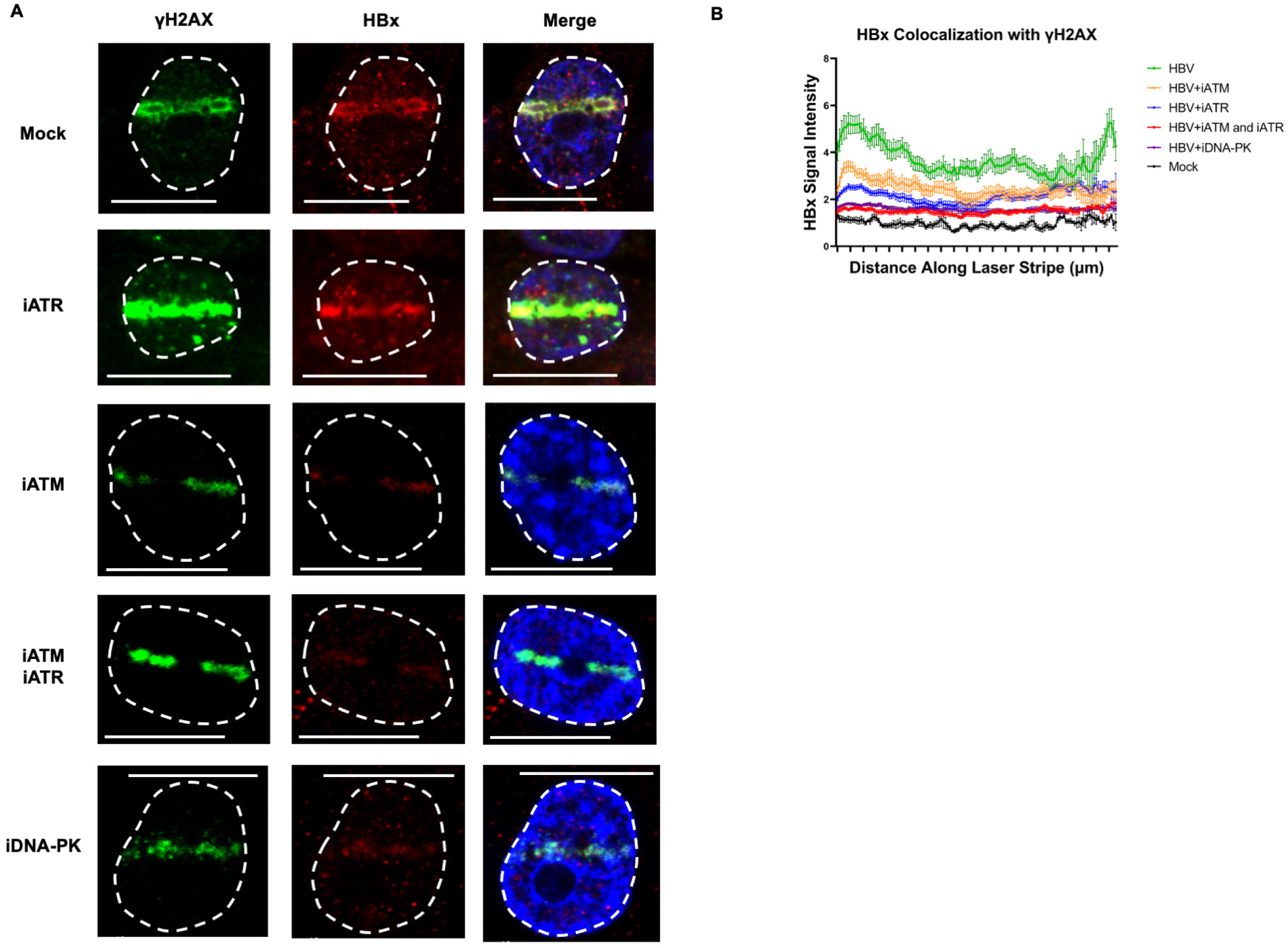
HBx proteins localize to cellular DNA breaks using host DDR kinase signaling. (A) Laser micro-irradiation assays were performed in HepG2-NTCP cells infected with HBV at 20 genome equivalents for 5 days before being processed for HBx protein (red) relocalization to cellular DDR (monitored by γH2AX staining; green). Images are representative of the independent inhibitor(s) each infected sample received 1 hour prior to micro-irradiation. (B) The signal intensity of HBx along the laser stripe was measured and normalized to overall signal intensity within the nuclei. The average intensity of HBx co-localizing with γH2AX between mock-treated cells and inhibitor-treated cells was averaged over at least 20 independent nuclei, represented as intensity profiles with the error bars at each point representing SEM.

## Discussion

In this study, we have dissected the connection between HBV infection and the cellular DNA damage response. HBV infection is sufficient to induce cellular replication stress early. However, this replication stress manifests as a cellular DNA damage response only upon conversion of rcDNA into cccDNA molecules 4-5 days post-infection and establishment of the latent viral reservoirs. The viral genome localizes to, and persists, in the vicinity of fragile genomic regions, presumably exacerbating replicative stress on the hepatocyte genome long-term. Importantly, the viral HBx protein localizes to induced cellular DDR sites in a PI3-kinase-like kinase dependent manner. This suggests that HBx and HBV genomes interact with cellular DDR regions by distinct mechanisms. Taken together, our results have begun to unravel the connection between HBV infection and cellular DNA damage signals that might regulate oncogenesis that leads to hepatocellular carcinomas.

Upon entry, DNA viruses localize to distinct nuclear sites that sustain the viral life cycle through gene expression, replication and persistence (70). These nuclear sites are rich in replication proteins, chromatin modifiers and transcription factors (9). These cytological observations have been collectively referred to as promyelocytic leukemia (PML) bodies, which also co-localize with HBV genomes and proteins during replication (71–73). Our findings in this study add to this catalog of HBV-associated cellular factors by including proteins in the DNA damage response and replication stress response pathways. However, it remains unclear whether DDR factors that colocalize with HBV are directly bound to the viral genome, associate with viral proteins or are bound to the cellular sites in the vicinity of the virus. While DDR proteins such as those in the ATR pathway are required to process rcDNA into cccDNA (31), it remains unclear whether they continue to persist on the viral genome long-term after conversion. Alternately, it remains conceivable that DDR proteins are the conduits for the recruitment of chromatin modifiers to the viral genome, facilitating the establishment of long-term latency, as has been observed for small DNA viruses such as AAV (74) and HPV (75, 76). Since AAV and HPV have been known to integrate into the host genome, it remains possible that DDR proteins co-localizing with HBV, as observed in this study, is a result of viral genomes integrating into the host. Interestingly, HBV integration events are at their highest at 5 dpi, correlating with the timepoint when they accrue maximum replication stress and DNA damage (77). The dissection of how and why DDR proteins associate with HBV, or the host genome in the vicinity of HBV components, warrants further study.

Studies by Shah and O’Shea in Adenoviruses revealed that virus and cellular genomes activate distinct DNA damage responses (78). In support of this model, we have previously discovered that the autonomous parvovirus MVM induces replication stress on the host genome that precedes induction of DNA damage by depleting the host of the single-stranded DNA binding protein RPA (57). The induction of replication stress by MVM occurs early in S phase and can be reversed by inhibiting the function of the replication fork helicase CDC7 (57). In contrast, our current study revealed that HBV-induced replication stress is observable immediately upon infection and is exacerbated over four to five days of infection, during which time HepG2-NTCP cells have undergone two rounds of mitosis (79). This suggests that HBV-induced replication stress on the host genome accrues progressively over time, eventually leading to the induction of cellular DNA damage. These observations resemble the replication stress phenotype induced by other tumor viruses such as HPV and mouse papillomavirus 1 (MmuPV1), that deploy the viral oncoproteins E6/E7 (13–15) and EBV that induces a hit and run mechanism using EBNA1 on the host genome at fragile sites (48).

Our observation that HBx oncoprotein relocalizes to induced cellular DDR sites in an ATM, ATR and DNA-PK dependent manner reveals novel features about the biology of HBV infection. Prior studies using parvovirus systems have shown that both viral genome and viral proteins localize to induced cellular DNA damage response sites. This localization is mediated by the non-structural protein NS1 (69). In contrast, however, the only DNA-binding protein encoded by HBV is the polymerase, which is presumably only used during reverse transcription (29). HBx on the other hand does not bind to the viral genome, instead perturbing the cellular machinery (80, 81). It has been shown that HBx triggers degradation of the cohesin subunit SMC5/6 (30). Doing so presumably contributes to accumulation of DNA damage. Interestingly SMC5/6 also restricts the transcriptional activity on HBV cccDNA (80, 81). Since cohesin proteins are also involved in maintenance of replication fork stability, we propose that HBx-triggered degradation of SMC5/6 impairs host replication forks that leads to the accumulation of cellular DNA damage. However, these mechanisms require further investigation in synchronous models of HBV infection that are currently unavailable.

Chromosome conformation capture technologies have proven to be invaluable tools to monitor where DNA viruses localize in the nucleus in an unbiased and high throughput manner. Prior studies that have deployed these techniques to investigate HBV genome localization have found that HBV genomes associate with heterochromatin sites on the human epigenome (51). We have previously used these technologies to show that parvoviruses localize to cellular DDR sites associated with transcriptionally active and accessible chromatin (47, 82). Similar studies with herpesviruses have found that EBV genomes localize to cellular enhancers that are enriched with the transcription factors ZN770, ZN121, PAX5, PRDM6 and IRF3 (83). The major drawback of these chromosome conformation capture assay systems has been their inability to distinguish between different forms of viral genomes, particularly those generated by DNA viruses in the nuclear compartment of dividing cells. We have resolved this issue in the current study by incorporating T5 exonuclease treatment into the chromosome conformation capture assay, allowing us to monitor the localization of cccDNA molecules exclusively. We have discovered that cccDNA molecules make up almost half of the total cellular sites that are measurably associated with HBV genomes. Since cccDNA molecules make up only 5-12 copies of DNA molecules per infected cell nucleus (54), these findings suggest most other genomic forms are either not tethered to the host genomic sites or converge on a small subset of genomic hotspots. Our assays combining in-silico prediction and imaging reveal that cccDNA molecules of HBV localize to cellular sites containing the DDR protein CHOP. Since CHOP has been implicated in oncogenic progression of hepatocytes leading to liver cancer (68), this might be one of the connecting links between HBV-induced DDR, cirrhosis and oncogenesis. Our Immuno-FISH assays corroborates the possible association between the HBV genome and CHOP, where we see sites of close localization between both. We surmise from these observations that by CHOP-mediated induction of reactive oxygen species (84) might exacerbate the replication stress in the host that are necessary for the conversion of its rcDNA to cccDNA. As a final consequence of this action, it is likely that CHOP overproduction causes cell death and inflammation which can stimulate tumor formation when dysregulated.

In conclusion, our work shows that HBV-induced replication stress begins at the onset of infection and persists over time and that this stress consequently induces the colocalization of several cellular DDR factors to HBV genomes and proteins. Furthermore, HBV cccDNA is preferentially located at cellular fragile sites, many of which are associated with important cellular transcription factors, such as CHOP. These findings open new avenues to investigate how HBV genomes and proteins interact with the host nuclear environment to cause oncogenic transformation.

## Materials and Methods

### Cell lines and viral stock preparation

Human hepatocyte cells HepG2 which overexpress the NTCP receptor [HepG2-NTCP; (85)] were maintained in Dulbecco’s modified Eagle’s medium/Nutrient Mixture F-12 (DMEM/F-12, high glucose; Gibco) supplemented with 5% Serum Plus (Sigma Aldrich) and 50 ug/ml penicillin/streptomycin (Gibco). HepG2-NTCP cells were maintained on collagen coated dishes for optimal growth and morphological characteristics. Tissue culture dishes were overlaid with 50 μg/ml rat tail collagen (BD Biosciences) in 0.02 N acetic acid solution and dried at room temperature for at least 1 hour for coating. The plates were then washed with phosphate-buffered saline (PBS) and used immediately or stored at 4 degrees Celsius overnight until use. For all imaging studies on cells plated on glass coverslips and laser micro-irradiation, collagen coating was used to aid in cell adherence. Cells were cultured in 5% CO2 at 37 degrees Celsius.

The Human hepatocyte cell line [HepAD38; (86)]was maintained in Dulbecco’s modified Eagle’s medium/Nutrient Mixture F-12 (DMEM/F-12, high glucose; Gibco) supplemented with 5% Serum Plus (Sigma Aldrich), 50 ug/ml penicillin/streptomycin (Gibco), 50 mg/ml gentamicin (Gibco) and 1 mg/ml doxycycline for the purposes of repression. HBV stock was produced by culturing HepAD38 cells until 70% confluency. Once confluent, doxycycline was removed, and cells were cultured in 5% CO2 at 37 degrees Celsius. Supernatant was collected every four days. Viral genome concentrations were calculated using qPCR compared with serial dilutions of known genome copies of the HBV infectious clone plasmid. HBV stock was maintained at -80 degrees Celsius.

### Viral infection

HepG2-NTCP cells were seeded into 6-well plates with DMEM/F-12 and incubated in 5% CO2 at 37 degrees Celsius for four hours. Fresh Spinoculum was made (DMEM/F-12 supplemented with 3% Serum Plus, 2% DMSO, 1% NEAA, and 4% PEG 8000). Solution was then filtered sterilized with 0.22um filter (Corning) Medium was switched with Spinoculum medium and HBV stock was added to 20GEQ. Plates were then wrapped with parafilm and then centrifuged at 1000xg at 37 degrees Celsius for 1 hour. Parafilm was removed and plates were incubated in 5% CO2 at 37 degrees Celsius for 5 days.

### Plasmids and transfections

All transfections were performed with linear polyethyleneimine (PEI) with a molecular weight (MW) of 25,000 (Polysciences). Cells were seeded the day before transfection in culture dishes pre-treated with collagen. Transfections were carried out in cells that were 60 to 70% confluent. The total mass of DNA transfected per condition was 1 μg/3.8 cm^2^ culture growth area and adjusted accordingly. The DNA to PEI ratio of 1:3 was used in NA-PEI-Opti-MEM mix. The media on transfected cells was replaced with fresh prewarmed medium between 6 to 18 hours post-transfection. The TMA153 plasmid referred to in this study as pHBV, is the same as the previously published pTMA153 and pLJ144, expressing full length HBV pgRNA, Cp, P, and all 3 envelope proteins (58). The pHBVΔHBeAg plasmid, previously referred to as pHC-9/3142, possesses a deletion in the cis-acting signal for pgRNA encapsidation, leading to the lack of production of the E antigen.

### Immunofluorescence imaging

Cells were fixed with 4% paraformaldehyde for 10 minutes at room temperature and then washed with PBS. 0.1% Triton X-100 was added for 10 minutes to permeabilize the cells. Samples were washed with PBS and blocked for 30 minutes with 3% BSA in PBS. Cells were then incubated at room temperature with the indicated primary antibodies for 1 hour, washed with PBS, and incubated for 30 minutes with the indicated secondary antibodies in 3% BSA. Coverslips were washed with PBS and mounted onto slides with Fluoromount containing DAPI (Southern Biotech).

### DNA fiber analysis

HepG2-NTCP cells were plated onto 6-well plates and infected by spinoculation and incubated for the indicated number of days. At the end of the indicated days, cells were pulsed with 20mM IdU in complete media for 20 minutes. Samples were washed with PBS and pulsed with 50mM CldU for 20 minutes in media. Cells were then pelleted for 5 minutes at 5000xg at room temperature and supernatant was discarded. Pellets were resuspended in 150ul of complete media and stored on ice. Two 2ul resuspensions were then pipetted onto the top of a positively charged slide at each end. 7ul of DNA Lysis buffer was added to each resuspension and pipetted gently four times. Samples were left at room temperature for 5 minutes on an even surface. Slides were then tilted at a gentle angle and left for 15 minutes for solutions to spread across the slide. While waiting, a solution of 3:1 methanol/acetic acid was made in a Coplin staining jar. Slides were added upright to the 3:1 solution and left for 5 minutes to allow the DNA to become fixed. Using a new jar, slides were washed three times in PBS and then denatured in a 2.5 M HCl solution for 1 hr at room temperature. Slides were washed three times with PBS. Laying the slides on a flat surface, 500ul of 3% BSA was added and left for 30 minutes. After blocking, the cells were stained with Abcam rat anti-BrdU (1:1000) and BD Biosciences mouse anti-BrdU (1:500) at room temperature for 30 minutes with a coverslip gently laid on the slide. After 30 minutes the coverslip was quickly removed, and the slides were washed with 0.1% Tween 20 in PBS three times. Samples were stained with anti-rat Alexa Fluor 488 and anti-mouse IgG1 Alexa Fluor 568 (1:1000) at room temperature for 30 minutes under covered conditions. Samples were washed with 0.1% Tween 20 in PBS 3 times and cover slips were affixed to slides using ProLong Gold Antifade Mountant (Thermo Scientific). Fibers were then imaged with a Leica Stellaris confocal microscope using a 63X oil immersion objective lens. Fiber lengths were measured using Digimizer software (MedCalc Software Ltd).

### Laser micro-irradiation

To perform laser micro irradiation, HepG2-NTCP cells were plated at 200,000 in three wells of a 6-well plate. The samples were then spinfected per protocol. After three days of incubation, the cells were gently trypsinized and centrifugated at 1,200 rpm before being transferred to collagen coated glass-bottomed dishes (MatTek Corp, Ashland, MA, USA) with complete media. After an additional two days of incubation the plates were supplemented with fresh complete media and 3ul of Hoechst dye (ThermoFisher Scientific, Madison, WI, USA) was added to sensitize the cells. Samples were then irradiated using a Leica Stellaris DMI8 confocal microscope using 63X oil objective and 2X digital zoom with a 405 nm laser using 25 percent power at 10 Hz frequency for 2 frames per field of view. Regions of interest (ROIs) were selected across the nucleus (edges of the nuclei were demarcated and visualized by Hoechst staining). Samples were then processed for immunofluorescence.

### Micro-irradiation imaging analysis

The intensity of HBx signal over the laser micro-irradiated region was quantified using the plot profile tool on FIJI software as we have previously established (69, 87). Signal intensities were measured at regular intervals from the left end to the right end of the region of interest (ROI) of the irradiated HepG2-NTCP cell nuclei. Values were averaged for 20–40 nuclei in 2–3 biological replicates at each position along the ROI. Quantification of the extent of HBx that relocalized to microirradiated sites relative to total HBx levels in the irradiated cell was computed by taking the following ratio: (HBx intensity along the microirradiated stripe)/(total HBx intensity in the nucleus). Images were processed with FIJI analytical software by outlining the ROI around the nucleus demarcated by the DAPI channel to measure total intensity of HBx while an additional ROI was drawn to outline the microirradiated stripe represented by the γH2AX channel.

### Fluorescence in-situ hybridization (FISH) assays

HepG2-NTCP cells were plated on coverslips and spinfected as per protocol. At five days post-infection, cells were fixed with 4% paraformaldehyde at room temperature for 10 minutes. The cells were then washed with cold PBS for 5 minutes and then immersed in 70% ethanol and permeabilized overnight at 4 degrees Celsius. Ethanol was aspirated and 10% formamide in 2X SSC buffer was added for 2 hours at room temperature to pre-denature the DNA. Amino-11-dUTP was conjugated to 20 basepair primers complementary to the target regions on the HBV and host genomes. NHS esters were used to conjugate fluorophores (AF-488 and AF-568, Thermo Scientific) to the ends of DNA primers. 2μL of each of the FISH probes were mixed in 40 μL of FISH hybridization buffer, mixed and added to the surface of the glass slide. The coverslip-containing cells were inverted over the FISH hybridization-probe solution and edges were sealed with rubber cement. Hybridization was carried out overnight at 37 degrees Celsius. Samples were washed twice in 2X SSC containing 0.1% Triton X-100 for 3 minutes each at 37 degrees Celsius. Samples were washed twice with 2X SSC at 37 degrees Celsius for 3 minutes each. Samples were mounted using DAPI-containing Fluoromount and imaged using a confocal microscope.

### Immuno-FISH assays

Immuno-FISH assays were performed on HepG2-NTCP cells that were cultured and infected as described for Immunofluorescense assays. 20 basepair DNA oligos that were complementary to the HBV genome (ACAAGGTAGGAGCTGGAGCA and GTAGGCTGCCTTCCTGACTG) were designed using Primer3 (88) and purchased from IDT. Oligos were pooled and labelled with 250 μM Amino-11-dUTP (Thermo Scientific) and TdT enzyme (Promega). Labelling reactions were carried out overnight at 37 degrees Celsius and were then terminated by incubating at 70 degrees Celsius for 10 minutes. Labelled oligos were precipitated in isopropanol, washed in 75% ethanol and dissolved in 15 μL of 0.1M sodium bicarbonate (pH 8.3). 0.75 μL of 20mM NHS esters (Thermo Scientific) were conjugated to the aminoallyl-tagged oligos for 2 hours in the dark. The labelled oligos were precipitated with isopropanol, washed in 75% ethanol twice and dissolved in 50 μL nuclease free water. The labelled oligos were precipitated in isopropanol, transferred to a PCR clean-up column (Promega, A9282) and centrifuged. Columns were washed twice in 80% pre-chilled ethanol and eluted in 50 μL nuclease free water. HBV infected HepG2-NTCP cells were CSK pre-extracted for 3 minutes in CSK buffer followed by CSK buffer containing Triton X-100. Samples were washed with PBS and fixed in 4% paraformaldehyde for 10 minutes at room temperature. Cells were permeabilized with permeabilization buffer for 10 minutes before being washed in PBS and denatured in 10% formamide solution in 2X SSC for 2 hours at 37 degrees Celsius. 2 μL of probe was mixed with 40 μL of FISH Hybridization buffer, mixed and added to the surface of the glass slide. The coverslip-containing cells were inverted over the FISH hybridization-probe solution and edges were sealed with rubber cement. Hybridization was carried out overnight at 37 degrees Celsius. Samples were washed twice in 2X SSC containing 0.1% Triton X-100 for 3 minutes each at 37 degrees Celsius. Samples were washed twice with 2X SSC at 37 degrees Celsius for 3 minutes each. Cells were then immunostained starting with blocking in 3% BSA in PBS as described above for immunofluorescence assays, mounted on DAPI-containing Fluoromount and imaged using a confocal microscope.

### Western blot

HepG2-NTCP cells were plated in 6-well plates at 200,000 cells. Samples were then spinfected with Spinoculum media at 20GEQ and left to incubate for 5 days. Cells were scrapped off and added to a microcentrifuge tube and centrifuged at 5,000 rpm for 2 minutes at room temperature. Samples were then resuspended in RIPA buffer and incubated on ice for 15 minutes. Samples were then centrifugated at 13,000 rpm for 10 minutes at 4 degrees Celsius. Supernatant was collected and the protein sample concentration was calculated using a bicinchoninic acid (BCA) assay (Bio-Rad).

### Viral chromosome conformation capture combined with T5 (V3C-T5-seq) assays

V3C-T5-seq assays were performed with BglII as the primary restriction enzyme to digest HBV-infected HepG2-NTCP chromatin. Briefly, samples were crosslinked in 1% formaldehyde for 10 minutes before being quenched in 0.125 M glycine on ice for 5 minutes. Cells were lysed in NP-40 lysis buffer for 10 minutes on ice, before resuspending the nuclei in restriction enzyme buffer (Cutsmart Buffer). Samples were permeabilized in 0.3% SDS for 1 hour on a 37 degrees Celsius shaker, sequestered in 2% Triton X-100 for 1 hour on a 37 degrees Celsius shaker. 1 ul of T5 Exonuclease (New England BioLabs) was added to the samples for 1 hr on a 37 degrees Celsius shaker. Samples were inactivated with 8 ul of 0.25 M EDTA before centrifuging at 5000 RPM for 5 minutes. Supernatant was removed and the pellet was resuspended in restriction enzyme buffer (Buffer 3.1) and digested overnight in 400 U of BglII enzyme with shaking. Samples were further digested with 300 U of BglII for 4 hours, inactivated at 65 degrees Celsius for 20 minutes in 1% SDS and sequestered with 1% Triton X-100 for 1 hour at 37 degrees Celsius. Chromatin was resuspended in 1.15X T4 DNA Ligase Reaction Buffer and intramolecular ligation was carried out at room temperature for 4 hours. Crosslinks were reversed and proteins digested with Proteinase K, RNAse A, and heat at 65 degrees Celsius. DNA was purified by phenol:chloroform:isoamyl alcohol extraction, isopropanol precipitation and using a PCR Purification kit (Qiagen). 3C DNA was eluted in 200 μL of DNA, secondary digested with NlaIII at 37 degrees Celsius overnight and circularized with 100 U of T4 DNA Ligase in 15 mL ligation reaction. The V3C-seq samples were precipitated by phenol:chloroform:isoamyl alcohol extraction, precipitated in isopropanol and resuspended in 100 μL of Buffer EB (Qiagen). Inverse PCRs were performed on the BglII-NlaIII fragments on the HBV genome using with inverse PCR primers tgccttctgacttctttccttcagt and cagtagctccaaattctttataaggg. DNA was diluted 1:100 in TE, before using them as templates for nested-inverse primer gacttctttccttcagtacg and tctttataagggtcgatgtc. The PCR reactions were pooled, purified using the PCR purification kit (Qiagen) and sequencing libraries were prepared using the NEB Ultra Kit. Twelve samples were pooled per run for paired end sequencing using an Illumina Next-seq 500 sequencer. Complexity of the sequencing reactions were increased by spiking in 25% phiX with the sequencing reactions.

### V3C-T5-seq analysis

High-throughput V3C-seq and V3C-T5-seq studies were aligned to the human hg38 reference genome using single end sequencing parameters in Bowtie2 alignment program (89). Samples were sorted with Samtools (90). Genome-wide coverage of the aligned intervals was measured with BEDtools (91). Independent biological replicates of the high-throughput sequencing studies were merged using BEDtools (91). Comparative analysis with published ChIP-seq and fragile site sequencing studies were performed using Deeptools package (92) on the Galaxy project server. The location of the V3C-T5-seq relative to that of published fragile site locations were computed by calculating the relative location of peaks in 2 Mb windows divided into 250 kb windows. V3C-T5-seq peaks containing more than 1000 reads in two independent biological replicates were selected for in-silico screening of transcription factor binding sites. BEDtools was used to obtain the DNA sequence of the intervals, which was used as the input for motif search using MEME. The transcription factor that associate with the 10 most common motifs were searched using FIMO and TOMTOM. Statistical significance of the V3C-seq with V3C-T5-seq analysis were performed using Jaccard analysis on BEDtools. The “observed” values were calculated using the intersection of all HBV sites with that of the cccDNA localization sites. This was compared with “permuted” values, where the HBV localization sites were intersected with a randomly generated set of genomic localization sites of the same size and number as cccDNA-associated regions.

### Data availability

All high-throughput sequencing data has been deposited in the Gene Expression Omnibus (GEO) repository and is publicly available under the accession number GSE261927.

**Table 1:** Table of antibodies used in this study.

| Reagent Type | Designation | Reference | Identifiers | Target species |
| --- | --- | --- | --- | --- |
| Antibody | γH2AX | EMD Millipore | 05–636 | Mouse |
| Antibody | Tubulin | EMD Millipore | 05-829 | Mouse |
| Antibody | Hepatitis B Core | OriGene | AP08118PU-N | Rabbit |
| Antibody | Hepatitis B Virus X | Abcam | ab39716 | Rabbit |
| Antibody | C/EBP-homologous protein (CHOP) | Cell Signaling | 2895 | Mouse |
| Antibody | ATRIP | Cell Signaling | 2737 | Rabbit |
| Antibody | BRCA1 | Invitrogen | MA1-23160 | Mouse |
| Antibody | PCNA | Abcam | ab29 | Mouse |
| Antibody | RPA70 | Abcam | ab176467 | Mouse |
| Antibody | Goat anti-Rabbit IgG, Alexa Fluor 488 | Invitrogen | A-11034 | Rabbit |
| Antibody | Goat anti-Mouse IgG, Alexa Fluor 568 | Invitrogen | A-11031 | Mouse |
| Antibody | Anti-rabbit IgG | Cell Signaling | 7074 | Rabbit |
| Antibody | Anti-mouse IgG | Cell Signaling | 7076 | Mouse |

## Acknowledgements

This research was funded partially by NIH/NIAID K99/R00 Pathway to Independence Award, grant number AI148511 to K.M. and The Wisconsin Partnership Program’s New Investigator Award (PERC Grant G-4942) to K.M. This project was supported in part by the American Cancer Society (ACS) grant IRG-19-146-54 to K.M. C.I.S.L. is funded by an NSF Graduate Research Fellowship Program award DGE-2137424. The author(s) thank the Precision Medicine Research Service of the UW Center for Human Genomics and Precision Medicine. We acknowledge Dr. Kavi Mehta (University of Wisconsin School of Veterinary Medicine) for guidance on replication stress studies, Dr. Andrew Huber (St. Jude Children’s Research Hospital) and Dr. Megan Spurgeon (University of Wisconsin School of Medicine and Public Health) for critical reading of the manuscript. We gratefully acknowledge Dr. Dan Loeb (University of Wisconsin School of Medicine and Public Health) for providing plasmids and cell lines critical for this study. Any opinions, findings, and conclusions or recommendations expressed in this material are those of the author(s) and do not necessarily reflect the views of the funding agencies. The authors declare no conflicts of interests.

## FIGURE LEGENDS

**Figure S1: HBV infection induces increased levels of γH2AX.** (A) Western blot analysis was performed at 5 dpi with uninfected, hydroxyurea induced, and HBV infected HepG2-NTCP cells. Western blots were imaged for double-stranded DNA breaks with γH2AX antibody and Tubulin as loading control. (B) Quantification of the signal intensities for γH2AX normalized to Tubulin in 2 independent biological replicates. The ratios were compared to that of HU-treated samples.

**Fig. S2: HBV localization to host cellular sites genome-wide.** Genome-wide occupancy of V3C-seq and V3C-T5-seq peaks in Hep-G2-NTCP cells infected with HBV at 5 dpi with the viewpoint on the HBV’s BglII restriction enzyme site.

